# Ruminosignatures associated with methane emissions and feed efficiency across geographies and cattle breeds

**DOI:** 10.64898/2026.02.19.706774

**Authors:** Ioanna-Theoni Vourlaki, Ori Furman, Ilma Tapio, Le Luo Guan, Sinéad M. Waters, David Kenny, Paul Smith, Stuart F. Kirwan, David Kelly, Ross Evans, Raquel Quintanilla, Antonio Reverter, Pâmela A. Alexandre, Fuyong Li, Philip C. Garnsworthy, Paolo Bani, Phillip B. Pope, Diego P. Morgavi, Itzhak Mizrahi, Yuliaxis Ramayo-Caldas

## Abstract

The cattle rumen microbiota represents a highly complex and dynamic ecosystem, whose organization and connection to host phenotypes are of the highest importance to food security and the environment. In this study, we analyzed the rumen microbiota, from 2,492 cattle belonging to five different breeds and production systems across five countries, categorizing them into microbial co-abundance groups referred to as Ruminosignatures. We identified twelve distinct Ruminosignatures, including two that were consistently observed across all populations and were dominated by the genus *Prevotella* and *UBA2810.* Additional Ruminosignatures showed breed-and diet-specific patterns and collectively explained 96–99% of the variance in rumen microbial composition. The abundances of several Ruminosignatures were associated with methane emissions and feed efficiency, and were influenced by host genetics, with heritability estimates ranging from 0.09 to 0.51. The Ruminosignature dominated by *UAB2810* was negatively associated with methane emissions across all datasets and positively linked to feed efficiency in Holstein from Italy and crossbred from Ireland. Additionally, the type of production system affects both the occurrence of Ruminosignatures and their impact on host phenotypes, emphasizing the need for context-specific approaches to modulate the rumen microbiome. Overall, our results offer new perspectives on the assembly of ruminal microbes and underscore the potential of the Ruminosignatures framework for microbiome-informed precision agriculture and breeding initiatives aimed at enhancing feed efficiency and minimizing the environmental impact of cattle farming.

## Introduction

Cattle production plays a central role in global food systems, with beef and dairy sectors contributing significantly to food security and economic sustainability [1]. However, similarly to other livestock species, cattle production faces global challenges related to the need to improve profitability and decrease its environmental impact. In this scenario, methane (CH₄) emissions and feed efficiency are critical traits that significantly impact both the productivity and environmental sustainability of cattle production [2, 3, 4].

Reductions in CH₄ emissions and improvements in feed efficiency can be achieved through a combination of management, nutritional strategies and selective breeding [5]. While management and nutritional strategies can have immediate effects, breeding provides a more permanent solution through genetic improvement. Although progress has been made in breeding for low methane emissions and high feed efficiency [7, 8], accurately measuring these traits in individual animals is complex and expensive, which limits the effectiveness of direct selection. Central to both aforementioned traits is the rumen microbiome, which mediates feed conversion and simultaneously produces CH_4_, via specific microbial taxa such as methanogenic archaea, protozoa, and associated bacteria. Integrating microbial signals that are linked to ruminal CH_4_ production into selection models offer an opportunity to enhance the accuracy of breeding programs [5, 9]. Most rumen microbiome studies have focused on specific cattle populations, such as European Aberdeen Angus × Limousin steers [10], Nordic Red dairy cows [11] or selected European and North American herds reviewed recently [5, 12]. This narrow scope limits the generalizability of findings across breeds and production systems, underscoring the need for multi-country datasets to assess microbiome potential and develop context-specific, microbiome-informed strategies.

Microbiome compositions are complex, sparse, and exhibit multivariate dependencies, making the analysis of these datasets particularly challenging. Instead of treating hundreds or thousands of individual microbial taxa as separate signals, the approach proposed by Frioux et al. [13] aims to reduce complexity and capture ecosystem-level dynamics by identifying microbial Co-abundance groups (CAGs). Microbial CAGs are defined as groups of microorganisms that display coordinated abundance patterns, often reflecting shared ecological roles and niches preferences, and consistently co-abundant behavior [14]. This approach has been demonstrated in humans [13] and pigs [15] and employs non-negative matrix factorization (NMF) to reduce dataset dimensionality by identifying latent co-abundance patterns that capture shared variance across taxa. Although traditional dimensionality reduction methods, such as principal component analysis (PCA) or ruminotypes, have been widely employed in studies of ruminant microbiota [16, 17, 18], they present certain limitations. PCA, although widely used, forces its components to be orthogonal and assigns both positive and negative loadings to taxa. This can mask biologically meaningful patterns and may combine taxa in ways that are not easy to interpret ecologically. Microbial CAGs, however, often exhibit overlapping or correlated activity patterns driven by ecological interactions. On the other hand, Ruminotype-based approaches classify samples into groups based on their dominant taxa, assigning each sample to one group only. Although these methods reduce dimensionality and can stratify samples to identify the taxon driving each cluster, they do not consider the ecological context. Consequently, the information they provide is limited in comparison. This study shows how we can use groups of rumen microbes that share patterns of bacterial abundance across individuals in diverse geographies and cattle populations as a variable connected to host traits such as methane emissions and feed efficiency traits.

## Material and Methods

### Study Populations and Experimental Design

The current study incorporates data from 2,492 animals from independent populations in five different countries that represent different cattle breeds and production systems. The discovery dataset was comprised from 818 cows: 409 Holstein lactating cows from United Kingdom (UK) and 409 from Italy [19]. A cohort consisting of 120 Holstein cows from France set [18] was used as independent validation. Two additional cohorts of beef and crossbreed cattle were employed as independent validation sets. The first one consisted of 705 multi-breeds (203 Angus, 114 Charolais, and 392 Kinsella composite hybrids) which were raised under standardized feedlot conditions at the University of Alberta’s Roy Berg Kinsella Research Ranch, Canada [20]. The second one includes 849 crossbreds from Ireland [21]. All animals were fed standardized diets appropriate for their production stage, with beef cattle receiving growth-phase-specific rations, crossbred from Ireland fed with a high energy concentrate-based diet [21, 22], and dairy cows maintained on total mixed rations for at least 14 days prior to sampling. Animal handling complied with the ethical standards and regulations established by Canada and the European Union directives on animal experimentation (European Commission, 2010). Ethical approval for the different studies can be found in the references cited above.

### Sample Collection and Processing

Rumen contents were collected using distinct but complementary protocols. For beef cattle, approximately 50 mL of rumen fluid was obtained via oro-gastric tubing before morning feeding at 293 ± 0.6 days of age, immediately frozen on dry ice, and stored at -80 °C (Li et al., 2019). Irish cattle sampled using the transesophageal rumen sampling device (FLORA rumen scoop; Guelph, ON, Canada). Dairy cow rumen samples were collected via esophagus using a Ruminator device (Profs Products, Wittybreut, Germany), with the first liter discarded to avoid saliva contamination, followed by collection of 500 mL of representative rumen fluid between 2-5 hours post-feeding [18, 19]. Samples were processed for immediate pH measurement, with aliquots preserved for volatile fatty acid following standard procedures.

### Phenotypic Measurements

Comprehensive production data were collected across the datasets (Figure 1). The beef and crossbred cattle trials recorded detailed feed efficiency metrics including dry matter intake (DMI), average daily gain (ADG), residual feed intake (RFI), and feed conversion ratio (FCR), CH_4_ emissions using Greenfeed monitoring system, Ireland only; and acetate and propionate levels were monitored for the Canadian beef cattle. The dairy cow trials monitored milk production parameters, body weight, and CH_4_ emissions using either breath sampling during milking (UK), Greenfeed (France, Italy). Table 1 presents a full description of the traits included in the association analysis while a summary of the datasets used across the analyses can be found on Supplementary Table 1.

**Figure 1.**
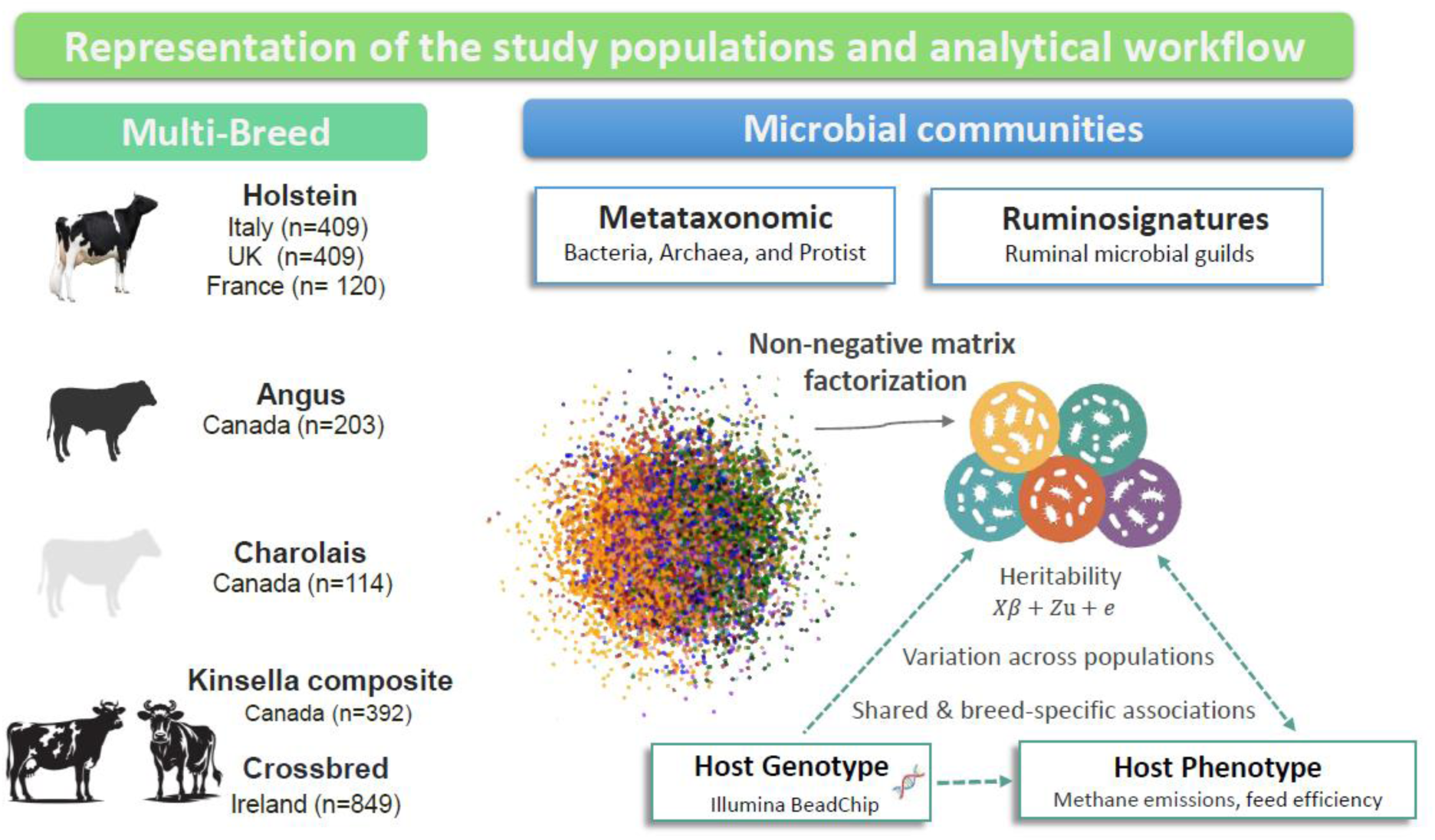
Summary of cattle breeds populations and analytical approach used in the study

**Table 1.** Summary statistics of the analyzed traits related to methane production, growth and feed efficiency across countries and genetic backgrounds.

| Trait | Country (Breed) | No samples | Mean (SD) |
| --- | --- | --- | --- |
| Methane production (CH<sub>4</sub> g/d) | United Kingdom (Holstein) | 407 | 430 (74) |
|  | Italy (Holstein) | 403 | 385 (106) |
|  | France (Holstein) | 65 | 506 (58) |
|  | Ireland (Crossbred) | 875 | 233 (51) |
| Methane production expressed as grams per kilogram of dry matter intake (CH<sub>4</sub> g/kg DMI) | United Kingdom (Holstein) | 402 | 19 (5) |
|  | Italy (Holstein) | 403 | 18 (5) |
|  | Ireland (Crossbred) | 883 | 20 (4.7) |
| Energy-corrected milk yield relative to dry matter intake (ECM kg/DMI) | United Kingdom (Holstein) | 402 | 1.5 (0.24) |
|  | Italy (Holstein) | 397 | 1.4 (0.33) |
| Average Daily Gain (ADG) | Ireland (Crossbred) | 779 | 1.5 (0.4) |
|  | Canada (Angus) | 183 | 1.3 (0.44) |
|  | Canada (Charolais) | 91 | 1.4 (0.33) |
|  | Canada (Kinsella composite) | 297 | 1.2 (0.35) |
| Feed Conversion Ratio (FCR) | Ireland (Crossbred) | 779 | 7.9 (2.3) |
|  | Canada (Angus) | 183 | 7.8 (2.5) |
|  | Canada (Charolais) | 91 | 7.8 (1.8) |
|  | Canada (Kinsella composite) | 297 | 7.6 (1.9) |
| Residual feed intake (RFI) | Canada (Angus) | 183 | 0.0 (0.6) |
|  | Canada (Charolais) | 91 | 0.0 (0.6) |
|  | Canada (Kinsella composite) | 297 | 0.0 (0.53) |
| Residual feed intake (calculated from NRC equations; RFI.NRC) | United Kingdom (Holstein) | 402 | -2.4 (3.24) |
|  | Italy (Holstein) | 397 | -0.5 (3.9) |
| Acetate-to-Propionate ratio (Ace: Prop) | Canada (Angus) | 200 | 3.5 (1) |
|  | Canada (Charolais) | 113 | 2 (0.7) |
|  | Canada (Kinsella composite) | 392 | 2.8 (0.9) |

### Microbial DNA extraction and sequencing

Total genomic DNA was extracted from rumen samples using optimized protocols for each study. The beef cattle samples underwent automated extraction using the QIAGEN BioSprint 96 workstation. Irish samples extracted using the DNeasy® PowerSoil® Pro Kit (QIAGEN GmbH) while dairy cattle samples employed a bead-beating protocol combined with the QIAamp DNA Stool Mini Kit (Qiagen). For bacterial and archaeal community profiling, the V1-V3 (UK, IT), V3-V5 (France), V4 (Ireland), and V6-V8 (Canada) regions of the 16S rRNA gene were amplified using established primer sets for bacteria and archaea. As described by Wallace et al. [19], the discovery dairy cohort additionally targeted protozoal 18S rRNA markers. All amplicons were generated on Illumina MiSeq platform.

### Bioinformatic Processing

Sequence data were processed through a standardized QIIME2 pipeline [24]. Raw reads underwent quality filtering (Phred score ≥20), adapter removal, and chimera detection. Amplicon sequence variants were generated at 99% similarity, with subsequent removal of low-abundance variants (<0.0001% total counts) and samples containing fewer than 8,000 reads. Taxonomic classification was performed using a customized GreenGenes2 (release 2024.09, [24]) reference database trained for the specific primer sets employed within each dataset, while the SILVA 138.2 database [25] was used specifically for taxonomic assignation of protozoan sequences.

### Identification of Ruminosignatures

Following the approach described by Frioux et al. [13], the existence of Ruminosignatures (RS) was assessed using NMF. NMF is an unsupervised dimensionality reduction technique that factorizes a non-negative input matrix into the product of two lower-rank matrices: ***W***, defined as the weight of the genera in each RS, and ***H***, which describe the RS abundance across samples. NMF identifies latent variables in the data capturing co-abundant patterns while performing a parts-based representation. Due to differences in primer sets across datasets, NMF was applied independently to each cohort. The optimal number of RS was accessed via a nine-fold bi-cross validation [26] as suggested in [13] for selecting the best rank in outer product models. For each of the nine folds, the model was trained on 8/9 of the data and evaluated on the remaining 1/9. This cross-validation procedure was repeated 200 times for a range of latent variables (k) from 2 to 30, with the input matrix randomly shuffled at each iteration. The most suitable number of RS was selected by monitoring the evaluation criteria associated with each k, including explained variance, and cosine similarity.

To perform bi-cross validation, we used the Python script **“*bicv_rank.py*”** from the repository of Frioux et al. [13]. The L1 regularization parameter (alpha) was estimated using the companion script **“*bicv_regu.py*”**, with the final value chosen according to the rank considered optimal. The corresponding alpha value for each dataset is provided in Supplementary Table 2. The analysis was carried out with the Python implementation proposed by Frioux et al. [13] in the same repository. The computations were run through the terminal in Python 3.9 [27], making use of SciPy v1.7.3 [28] and Scikit-learn v0.24.1 [29]. The analysis was performed on normalized genus-level relative abundance tables. Normalization was carried out by dividing the counts in each column (representing an individual sample) by the total sum of counts in that column, so that each column summed to 1 and all values were positive. Each configuration was executed for 100 iterations, with a maximum of 2000 updates, random initialization, a defined regularization ratio, the multiplicative update algorithm, and the Kullback–Leibler divergence as the β-loss function. Regularization was applied simultaneously to both the ***W*** and ***H*** matrices. We investigated the correlation patterns between RS and the abundance of methanogenic archaea. Particularly, for the Holstein discovery dataset, links with the abundance of protozoal ruminal communities was also explored.

### Association of Ruminosignatures with methane emission and feed efficiency

The association analyses were performed using a univariate linear model approach, applied separately to each of the datasets included in this study. All models were fitted in R [30] using the *lm* function, and the results were visualized with the *ggplot2* [31] package. Given the differences across datasets in both structure and available metadata, dataset-specific covariates were included in the models as described below. P-values < 0.05 were considered significant.

In the Holstein discovery dataset, we assessed the links between RS abundance and CH_4_ and feed efficiency-related traits, including CH_4_ yield per day (CH₄ g/d), CH_4_ yield per kilogram dry matter intake (CH₄ g/kg DMI), energy-corrected milk to DMI ratio (ECM kg/DMI), and residual feed intake based on NRC (RFI.NRC). Analyses were performed by country: Italy (IT) and the UK, incorporating the first five principal components to control for population structure. Farm of origin was included as a fixed effect (IT three levels and UK two levels). In the French validation dataset, RS associations with CH₄ g/d were assessed in the subset 65 cows with available measurements. Since CH₄ were not directly measured in the Canadian dataset, we used the acetate-to-propionate ratio (Ace: Prop) as a proxy for CH_4_ production. RS abundance was tested for association with

Ace: Prop and additional performance traits including ADG, RFI, and FCR. Analyses were conducted separately by breed: Angus (ANG), Charolais (CHAR), and Kinsella composite hybrid (HYB). All traits were corrected for age, while animal type (three levels: bulls, N = 70; heifers, N = 347; steers, N = 288) and diet (four levels) were included as fixed effects. Finally, for crossbred animals from Ireland, associations between RS abundance and FCR, ADG, CH₄ g/d, and CH₄ g/kg DMI were evaluated. Models included age and the first five principal components as covariates, while contemporary group, and animal type (three levels: heifer = 352; steer = 402; young bull = 130) were included as fixed effects.

### Host genotype and heritability estimation

Cows from the Holstein discovery dataset were genotyped with the Bovine GGP HD chip v2 that included 138,892 SNP, the Irish crossbred were genotyped with a custom chip (International Dairy and Beef Chip Illumina BeadChip), latter imputed to a 50K SNPs using Fimpute [32], while the other populations were genotyped with the Illumina BovineSNP50 v2 Genotyping BeadChip which contained 54,609 SNP (San Diego, CA, USA). Plink software [33] was used to remove SNPs that had a minor allele frequency less than 5%, that had more than 10% missing genotype data, mapping to the sex chromosome, or that did not map to the ARS-UCD1.3 bovine assembly.

The potential host genetic control on RS abundance was explored by estimating the genetic variance components. Specifically, for each of the datasets, the narrow-sense heritability (ĥ^2^) was estimated by implementing the Reproducing Kernel Hilbert Space (RKHS, [34]) model from the BGLR package [35] applied separately to each cohort. Prior to analysis, the abundance of each RS was subjected to an inverse normal (rank-based) transformation to ensure homoscedasticity in the model residuals.

After quality control the genotypic data for the Holstein discovery set included 109,193 SNP across 771 animals. The model was implemented jointly for both countries (IT and UK), with farm included as a fixed effect as follows:

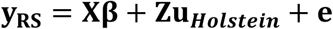

where 𝐲 is the vector of individual RS abundances, ***β*** is the vector of the systematic effects of farm (five levels) and 𝐗 its corresponding incidence matrix, 𝑢 is the vector of additive genetic random effects, whereas 𝐞 is the vector of residuals.

The Canadian dataset comprised genotypic information for 43,123 SNP across 685 samples, while the Irish dataset included 41,335 SNPs from 843 individuals. The mixed model was applied independently to each dataset, as described below:

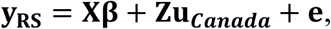

where, β is the vector of the fixed effects of animal-type (three levels), breed (three levels), diet (four levels), and 𝐗 their corresponding incidence matrix.

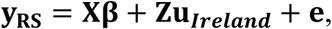

where, β is the vector of the covariate age, contemporary group and systematic effects of animal type (three levels) and dataset (two levels).

In all datasets random genetic effects were assumed to follow a multivariate normal distribution 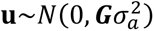, where 𝐆 is the genomic relationship matrix (GRM) calculated using VanRaden method and 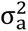 is the additive genetic variance. The GRM was obtained as 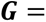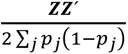 is the minor allele frequency of SNP *j,* and *Z* is the MAF-adjusted genotype matrix generated with the AGHMatrix package [36, 37]. For the estimation of genetic variances with BGLR, default priors were used. Thus, the narrow-sense heritability is defined as the proportion of RS variance explained by additive genetic effects as follows:

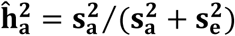

where 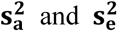 are the posterior estimates of the additive genetic and residual variances respectively. **BGLR** was run for 100,000 iterations with a burn-in of 500 and thinning interval of five. Convergence was assessed visually, and the corresponding plots are provided in Supplementary Figures 1–3 for each dataset. Posterior Gibbs samples representing 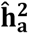, were entered into a Monte Carlo Markov Chain (MCMC) object using the as.mcmc() function from the coda package in R [38] while the 95% highest posterior density intervals (HPD95) were computed for each heritability estimate.

## Results

### Ruminosignatures profiles across Holstein populations

NMF analysis revealed the existence of six RS in the Holstein discovery population. The optimal number of RS (k = 6) was determined based on the plateau observed in both cosine similarity and explained variance metrics (Supplementary Figure 4), beyond which additional factors yielded negligible improvement. Notably, this model explained 99% of the variance and achieved a cosine similarity of 99% from the original genus abundance table (Supplementary Table 3).

The driving genera was defined as those with the highest contributions within each microbial CAGs. Following this criterion, the taxonomic composition of the six RSs was driven by *Prevotella* (RS-Prev), *UBA2810* (RS-UBA2), *Fibrobacter* (RS-Fibr), *RF16* (RS-RF16), *Treponema* (RS-Trep), and *Succiniclasticum* (RS-Succ). Figure 2 illustrates the composition of the six RSs and their corresponding genus contribution, displaying only genera with a relative contribution greater than 1% in at least one RS. As shown, RS profiles vary in both taxonomic structure and contribution. Although RS-Prev and RS-UBA2 are largely dominated by a few genera, the others (RS-Fibr, RS-RF16, RS-Trep and RS-Succ) represent microbial CAGs composed of multiple genera with low to moderate contributions. To validate the initial findings, we applied the same NMF-based approach to an independent Holstein dataset from France. In this cohort, the optimal rank was determined to be k = 7 (Supplementary Figure 5), capturing 99% of the variance in bacterial genus abundances and achieving a cosine similarity of 99% (Supplementary Table 3). Four RS including RS-Prev, RS-UBA2, RS-Fibr, and RS-RF16 were consistent with those identified in the discovery dataset. The remaining three were specific to the French population and dominated by: *Sodaphilus* (RS-Soda), *Succinivibrio* (RS-Succ), and *Butyrivibrio_A_168226* (RS-Buty) genera. The genus-level composition of each RS is shown in Supplementary Figure 6, with the dominant genera contributing between 14% and 74% of the total abundance within their respective microbial CAGs. In agreement with the results from the discovery dataset, RS-Prev and RS-UBA2 were dominated by a few genera, whereas the others represented microbial CAGs composed of multiple genera with low to moderate contributions (Supplementary Figure 6).

**Figure 2.**
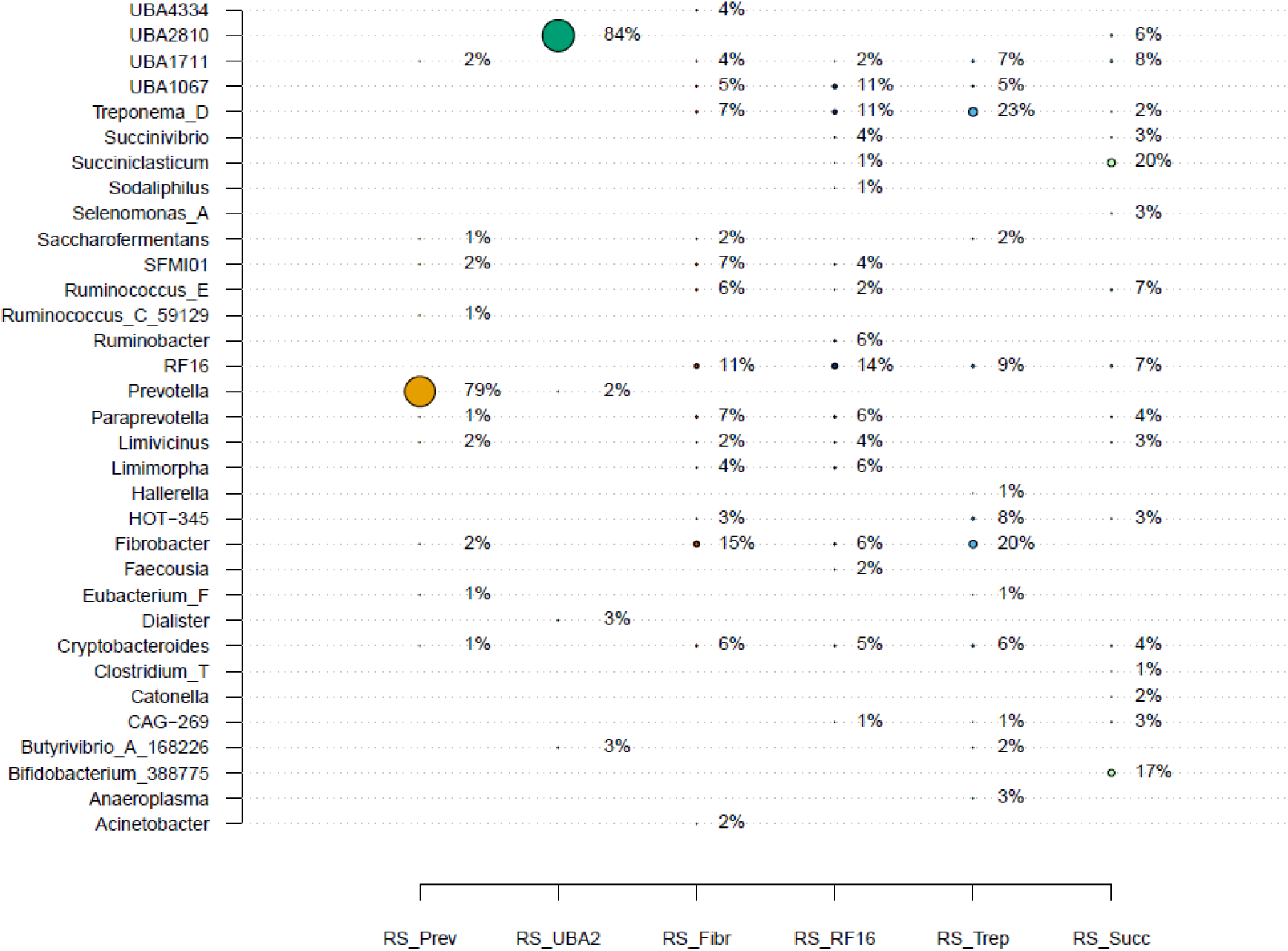
Relative genera composition of Ruminosignatures in the Holstein discovery set. Only genera with a contribution higher than 1% in at least one of the Ruminosignatures are shown.

### Ruminosignatures profiles across Canadian and Irish populations

Results from the Canadian cohort suggested the existence of eight RS (Supplementary Figure 7), collectively explaining 96% of the total variance in genus-level abundance data and achieving a cosine similarity of 98% (Supplementary Table 3). The main RS driving variation in this dataset were RS-Prev, RS-UBA2, RS-Rumi, RS-Succ, RS-SMFI, RS-Limi, RS-Soda, and RS-Cryp, corresponding to the dominant genera *Prevotella, UBA2810, Ruminococcus, Succiniclasticum, SMFI, Limimorpha, Sodaliphilus,* and *Cryptobacteroides*, respectively (Supplementary Figure 8). As shown in Supplementary Figure 8, and consistent with observations from Holstein populations, RS-Prev and RS-UBA2 were again primarily driven by their dominant genera, contributing 86% and 93% of the respective RS composition. Within the Irish crossbreed population seven RS (Supplementary Figure 9) were identified accounting for 97% of the total variance in the original genera matrix and capturing 98% of the cosine similarity (Supplementary Table 3). The taxonomic composition of the main RS was dominated by RS-Prev, RS-UBA2, RS-Rumi, RS-Succ, RS-SMFI, RS-RF16, and *Sharpea* (RS-Shar). As observed in other populations, RS-Prev and RS-UBA2 were primarily driven by their respective dominant genus, whereas the remaining RS consisted of microbial CAGs composed of genera with low to moderate contributions (Supplementary Figure 10).

### Population-level variation in Ruminosignature profiles

The comparison across all datasets revealed two RSs that were common across geographies, breed and populations: RS-Prev and RS-UBA2 (Figure 3). Another two (driven by *RF16* and *Succiniclasticum*) were common across three of the four datasets, while RS-Fibr, RS-SMFI, RS-Rumi, and RS-Soda were detected in at least two populations. The remaining RS appeared to be diet, breed or production system specific. RS-Trep was exclusive to the Holstein discovery dataset, while the RS driven by *Sharpea* was observed only in the Irish crossbreed population. RSs drived by *Limimorpha* and *Cryptobacteroides* were unique to the Canadian cohort, and *Succinivibrio* and *Butyrivibrio* were exclusively found in the French Holstein population (Suppelmentary Figure 6). When comparing Canadian beef cattle with the Irish cohort and Holstein discovery dataset, three common RSs were identified, dominated by *Prevotella*, *UBA2810*, and *Succiniclasticum*. In contrast, comparison with the French Holstein population revealed a different overlap, for the shared RS dominated by *Sodaliphilus*.

**Figure 3.**
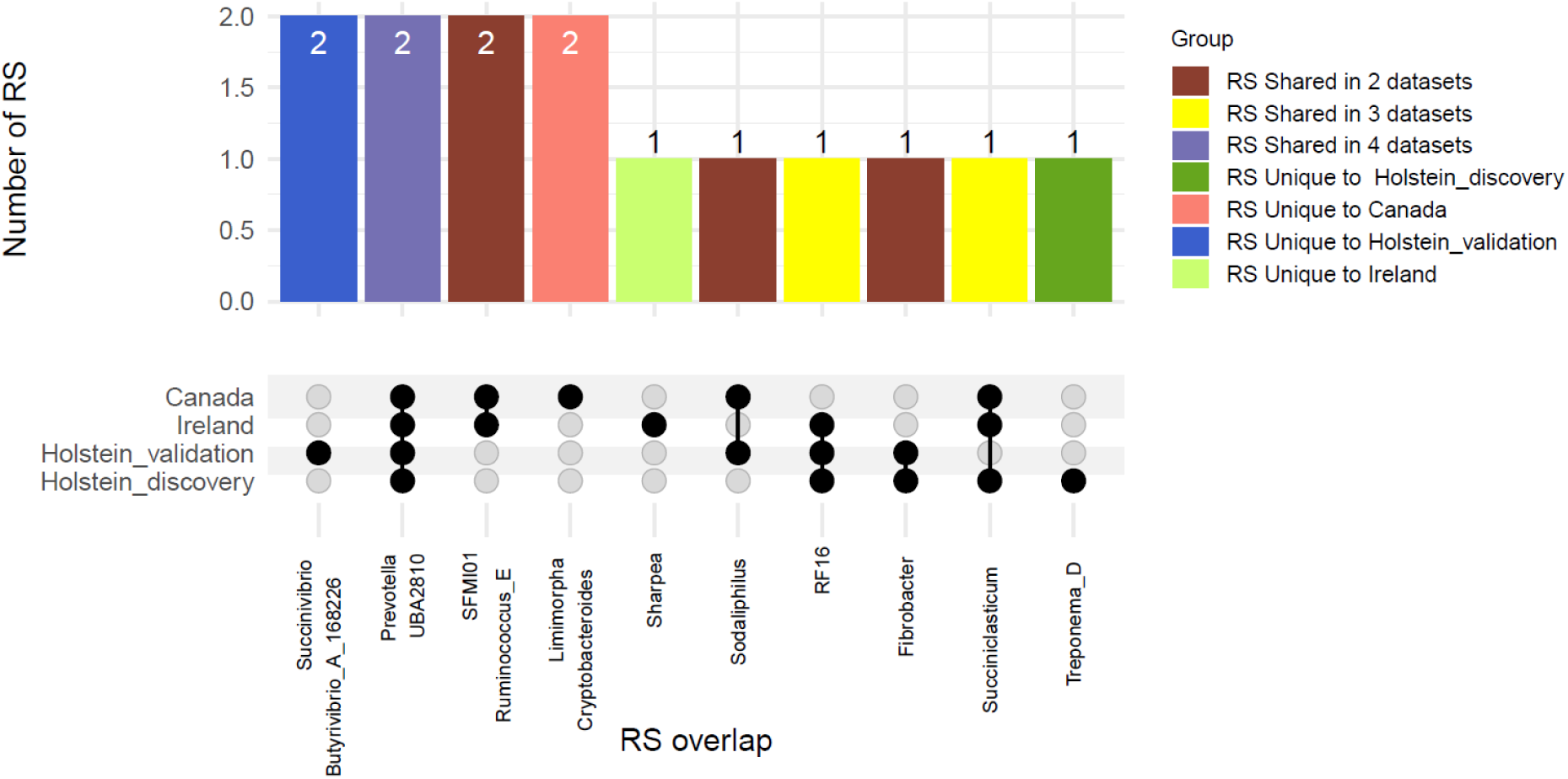
Comparison of Ruminosignatures intersection across all the datasets. The bar plot (top) shows the number of Ruminosignatures that are unique to each dataset or shared across 2, 3, or all 4 datasets, with colours indicating the grouping categories. The UpSet plot (bottom) displays the overlap structure of individual Ruminosignatures across the Canada, Ireland, Holstein-discovery, and Holstein-validation datasets. Filled circles indicate the presence of a given Ruminosignatures in each dataset. Together, the figure summarises both dataset-specific and commons Ruminosignatures across the study cohorts.

### Comparative analysis of taxonomy contribution across core Ruminosignatures

To gain deeper insight into RS assembly and the putative ecological roles of individual genera within the core microbial CAGs (RS-Prev and RS-UBA2), we performed a stratified analysis by dataset. For each dataset, genera showing a non-zero contribution in at least one of the RS were retained. Then, the different datasets were intersected, and the genera shared across all of them were identified. Figure 4 illustrates the fourteen genera commonly associated with at least one of the core CAGs across all cattle populations, revealing distinct yet recurrent patterns. As previously described, the core-RS were consistently dominated by *Prevotella* and *UBA2810*, that contributed the largest proportion of the RS community structure. Remarkably, despite differences in diet, genetic background, farm of origin, country or production system, the identity of the dominant genus remained consistent across all datasets. However, the relative contribution of the dominant genus varied in magnitude; for instance, *UBA2810* exhibited a higher contribution in the Canadian beef cohort (90%) and a lower (60%) in the Irish crossbred. Beyond the dominant genera, we also observed consistent trends among less prevalent genera that contribute to the composition of RS-Prev and RS-UBA2. For example, *Cryptobacteroides* appeared consistently linked to RS-Prev, with comparable contributions across datasets. Similarly, *Paraprevotella* exhibited strong co-occurrence patterns to RS-Prev across datasets. Notably, RF16 was detected exclusively in RS-Prev, whereas genera such as *Ruminococcus_E* and *Succinivibrio* displayed a marked shift in abundance depending on the RS type. CAG-269 followed a similar pattern in both RSs, although its highest contribution was observed in RS-UBA2. Overall, RS-Prev is robustly defined by *Prevotella*, while the presence and abundance of co-occurring genera appear to be context-dependent. Likewise, RS-UBA2 was consistently drived by *UBA2810*, though the composition of associated genera varies considerably between populations. While RS type remains a major driver of genus-level structure, population-specific factors also modulate the contribution of individual genera. RS-Prev and RS-UBA2 harbor distinct microbial signatures: some genera are consistently enriched in one RS across all datasets, serving as robust markers of RS identity, whereas others exhibit variable patterns, likely influenced by host or environmental context.

**Figure 4.**
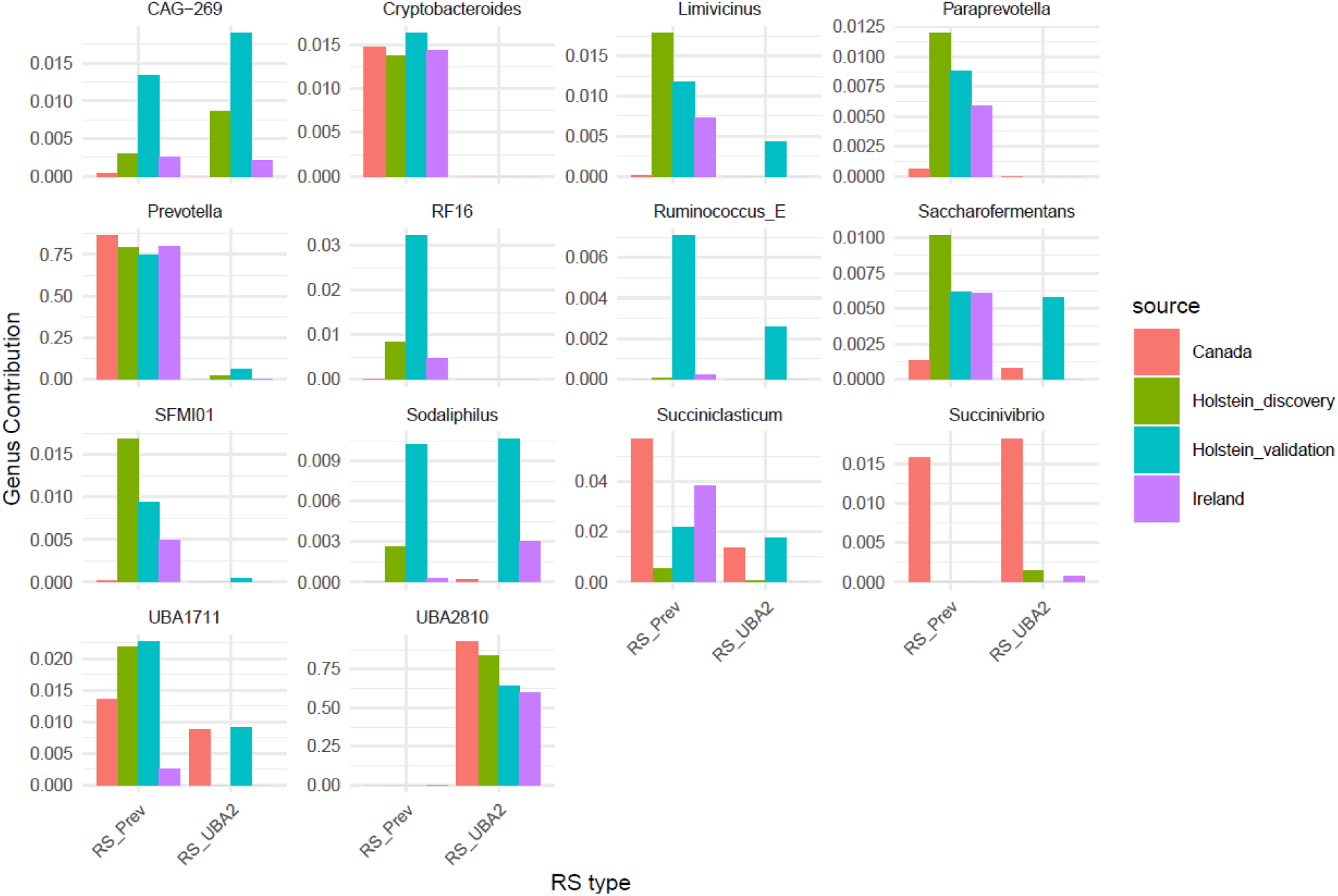
The bar plots panel illustrates the contribution of fourteen common genera identified across the four datasets, with distributions shown separately for the two core microbial CAGs, RS_Prev and RS_UBA2. The figure highlights the stability of dominant genera (e.g., Prevotella in RS_Prev, UBA2810 in RS_UBA2) alongside variation in the co-occurring taxa across datasets.

### Association with methane emissions and feed efficiency

Beyond characterizing the number and taxonomic composition of the RSs, we investigated their associations with CH₄ emissions and feed efficiency traits. To this end, linear models correcting by confounders (see Methods) were conducted to explore the relationship between the RS abundance and CH₄ and feed efficiency-related parameters. Remarkably, RS-UBA2 was consistently negatively associated (p < 0.05) with CH₄ emission across all breeds in each of the analyzed countries (France, Ireland, Italy, and the UK), and with Ace: Prop in Canadian beef populations (Figure 5). Conversely, RS-Prev showed a negative association with CH_4_ in Holstein animals but not in the Irish crossbred population (Supplementary Figure 11). On the other hand, RS-RF16 generally exhibited positive associations with CH₄ emissions, except in the Canadian population, where the Ace: Prop was employed as proxy of CH_4_ emission (Supplementary Figure 11). In the same dataset from Canada, RS-SMFI and RS-Rumi were significantly (p < 0.05) positively associated with a lower Ace: Prop. Although the Canadian animals were fed rumensin and the model accounts for dietary differences, the specific effects of rumensin on the ruminal microbiota remain unclear. Further research is needed to determine whether it alters community composition or the abundance of particular microorganisms.

**Figure 5.**
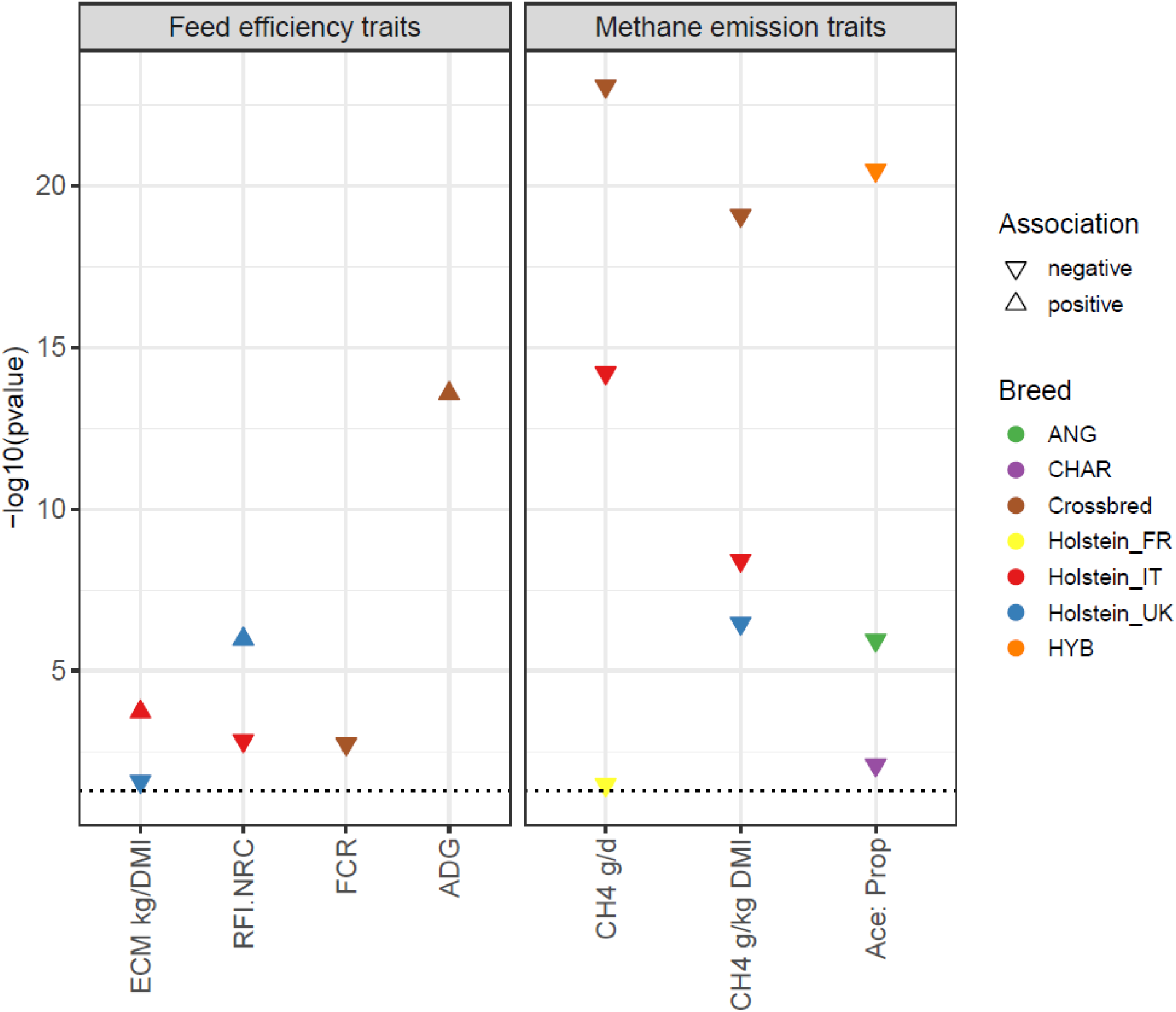
Linear associations between RS-UBA2 abundances and methane or host performance traits. Arrows indicate the direction of the relationship between the Ruminosignature driven by UBA2 and host trait. Only significant associations are shown, although the traits were measured in additional breeds not displayed in the plot.

We also observed inverse correlation patterns suggesting co-exclusion among microbial CAGs associated with CH_4_. For example, in the Holstein populations, RS-UBA2 showed a negative correlation with the abundance of RS-RF16 (R = -0.65, P < 0.001). RS-UBA2 was also negatively correlated (R = -0.39, P < 0.001) with the abundance of the methanogenic archaea *Methanobrevibacter*, while the abundance of RS-RF16 was positively correlated with *Methanobrevibacter* (R = 0.37, P < 0.001). In the Irish cohort, RS-UBA2 negatively correlated with the abundance of RS-Rumi (R = -0.42, P < 0.001) and negatively with *Methanobrevibacter* (R = -0.25, P < 0.01), whereas the abundance of RS-Rumi positively correlated with *Methanobrevibacter* (R = 0.49, P < 0.001). Interesting, this same pattern, (RS-UBA2 vs RS-Rumi = -0.24, P < 0.01; RS-Rumi vs *Methanobrevibacter* = 0.32; P < 0.001) was confirmed in the Canadian dataset.

Regarding feed efficiency traits, RS-UBA2 was negatively associated with both FCR and RFI and positively associated with ADG in the crossbred Irish population (Figure 5). This indicates that crossbreed animals from Ireland with higher abundance of RS-UBA2 tend to utilize feed more efficiently resulting in higher daily growth rates while emitting less CH₄. A similar association pattern was observed in Holstein cattle from Italy, where RS-UBA2 showed a positive association with energy-corrected milk per dry matter intake (ECM kg/DMI) and a negative association with RFI. In contrast, Holstein cows from UK exhibited the opposite trend, suggesting that the relationship between RS-UBA2 and feed efficiency in Holstein cows may be influenced by population-specific factors such as diet, environment, or management practices.

### Host genetic control on Ruminosignatures abundance

Table 2 shows the posterior mean estimates of heritability 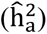 for the abundance of each RS across the beef (Canada), beef-dairy crossbred (Ireland), and Holstein (UK and Italy) datasets. France validation sets were not included in this analysis because of the small number of available animals. When comparing heritability estimates between common and dataset-specific RSs, the highest values were observed in the Irish crossbred dataset, for which the abundance of RSs driven by *UBA2810*, *SMFI*, *Ruminococcus_E*, and *Succiniclasticum* showed posterior mean heritability of 0.48, 0.49, 0.51, and 0.58, respectively. Additionally, the RS driven by *Sharpea* exhibited a moderate heritability estimate of 0.38 in this crossbred population. The Holstein discovery dataset showed intermediate heritability values for the different RS abundances 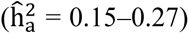, while the Canadian dataset displayed the lowest estimates 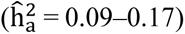. Importantly, in all cases, the posterior mean and the 95% highest posterior density interval did not include zero, supporting the robustness of these estimates (Table 2).

**Table 2.** Summary of posterior mean of heritability and 95% highest posterior density interval (HPD95) for each ruminosigantures across datasets.

| RS | Country | $\hat{h}_a^2$ | HPD95 [lower upper] |
| --- | --- | --- | --- |
| RS_Prev | Canada | 0.11 | [0.04 0.19] |
|  | Holstein | 0.17 | [0.1 0.25] |
|  | Ireland | 0.19 | [0.07 0.32] |
| RS_UBA2 | Canada | 0.14 | [0.5 0.24] |
|  | Holstein | 0.27 | [0.15 0.39] |
|  | Ireland | 0.48 | [0.33 0.63] |
| RS_SMFI | Canada | 0.17 | [0.09 0.29] |
|  | Ireland | 0.59 | [0.45 0.72] |
| RS_Rumi | Canada | 0.13 | [0.06 0.22] |
|  | Ireland | 0.49 | [0.34 0.65] |
| RS_Succ | Canada | 0.15 | [0.08 0.25] |
|  | Holstein | 0.15 | [0.1 0.22] |
|  | Ireland | 0.51 | [0.36 0.65] |
| RS_Limi | Canada | 0.13 | [0.06 0.21] |
| RS_Soda | Canada | 0.12 | [0.05 0.21] |
| RS_Cryp | Canada | 0.09 | [0.04 0.16] |
| RS_Fibr | Holstein | 0.25 | [0.15 0.35] |
| RS_RF16 | Holstein | 0.24 | [0.13 0.36] |
|  | Ireland | 0.25 | [0.11 0.39] |
| RS_Trep | Holstein | 0.27 | [0.15 0.38] |
| RS_Shar | Ireland | 0.38 | [0.22 0.55] |

## Discussion

In this study, we proposed the term Ruminosignatures to define the existence of microbial CAGs within bovine ruminal microbiota. The identification of microbial CAGs through NMF represents a significant methodological advance in the study of microbial ecosystems. Unlike traditional dimensionality reduction approaches such as principal component analysis or ruminotypes the NMF-based decomposition provides a parts-based representation that is biologically interpretable and ecologically relevant. PCA imposes orthogonality constraints among components [39], which may obscure overlapping ecological processes that occur simultaneously in the rumen. In contrast, NMF enables taxa to contribute to multiple latent components, capturing co-abundance patterns that reflect coordinated ecological behavior [13]. The advantage of NMF is that these latent variables can reveal underlying biological processes in a more interpretable way. Samples can be expressed as additive mixtures of multiple communities, reflecting the fact that microbial CAGs often interact rather than exist in isolation. Moreover, unlike PCA, NMF does not require variables to be orthogonal or independent. Instead, it searches for latent co-abundance patterns that capture shared variance across taxa. Each microbial CAGs represents a distinct co-abundance pattern, yet taxa can overlap and covary across samples. Additionally, this approach provides comprehensive ecological information, detailing which features are co-abundant within each RS and the relative abundance of each RS within individual samples. This approach mirrors the additive nature of microbial ecosystems, where communities interact and respond collectively to environmental and host-specific factors [13]. On the other hand, NMF-ecological decomposition contrasts sharply with ruminotypes, where samples are assigned to discrete clusters. While ruminotype can identify dominant community structures, they do not account for overlapping microbial memberships or continuous variation in community composition. In contrast, NMF quantifies the contribution of each RS to every sample, enabling a more nuanced description of the rumen microbiome continuum.

This analysis reveals that the cattle ruminal microbiota can be accurately described by combinations of at least 12 RS, which, depending on the cattle population analyzed, capture between 96% and 99% of the variance in the bacterial community. This study demonstrated the heritable nature of RS abundance 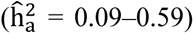, supporting a significant influence of host genetics on bacterial community assembly, which aligns with previous reports of low to moderate heritability for core microbiota as well as for some rumen taxa [19, 20, 40, 41, 42].

Suggesting their fundamental role in the microbial ruminal ecosystem, two core microbial RS dominated by *Prevotella* and *UBA2810* genera were consistently identified across 2,492 diverse cattle breeds from five countries. The existence of breed, diet, and production-system specific RSs was also observed, reinforcing the influence of diet, production system, and host genetics on the assembly of the ruminal microbiota [20, 43]. For example, RS-SMFI was exclusive to beef cattle, while RS-RF16 was specific to Holstein or crossbred populations. Intriguingly, the RS profiles observed in the Irish crossbred population exhibited the greatest overlap (six out seven) with those identified in both beef and dairy cattle. This finding reflects the diverse genetic background of the Irish crossbred population and highlights the potential of the NMF-based approach to identify both commons and breed-specific microbial CAGs.

Likewise, it was described that breed-specific and across-breed associations between RS abundance and key traits such as growth performance, feed efficiency and CH_4_ emission. RS-RF16 displayed a consistent positive association with CH₄ in Holstein cows from France, Italy, and the UK. Among the breed-specific, the RS driven by *Sharpea* detected in the Irish crossbred population showed a negative link with CH₄. *Sharpea* has been proposed as keystone genera driving a ruminotype *“S”* negatively associated with CH₄ yields in sheep [17] and was also suggested as a biomarker for low CH_4_ production in cattle [21]. The increased abundance of *Sharpea* in the rumen promotes a fermentation pathway that favors lactate and butyrate production. This shift reduces the availability of hydrogen for methanogens, thereby leading to decreased methane emissions in ruminants [44, 45]. In the Irish crossbred and the French Holstein populations, a positive association between the RS driven by *Ruminococcus* and CH₄ emissions was observed. In contrast to *Sharpea*, an elevated abundance of *Ruminococcus* has been documented in a ruminotype *’H’*, which was linked to higher CH₄ emissions [17]. These findings demonstrate that the RSs models confirm key ecological roles known from previous ruminotype concepts, while enabling the detection of gradual shifts within the cattle ruminal bacterial community.

Recently, Tovar-Herrera et al. [46] proposed 14 genera as the core of the ruminant microbiome. These core taxa are characterized by broad metabolic potential, nutritional independence, and extensive redundancy across key biosynthetic pathways. Such functional traits, together with their ability to degrade plant polysaccharides through diverse glycosyl hydrolases and to synthesize essential metabolites such as amino acids and vitamins, highlight their ecological role as keystone taxa within the ruminal ecosystem and lead to the suggestion that these taxa play a central role in supporting host metabolism and ecosystem function [46]. In the present study, we identify Ruminosignatures that consistently co-vary in abundance across animals, without a priori assumptions regarding function, while nonetheless showing consistent associations with host traits. Notably, seven of the core genera reported by Tovar-Herrera et al. [46] were among the major drivers of the Ruminosignatures (*Prevotella, Fibrobacter, Succiniclasticum, Ruminococcus, Treponema, Succinivibrio*, and *Butyrivibrio*). Moreover, as illustrated in Figure 4, five of them (*Saccharofermentans, Succiniclasticum, Succinivibrio, Ruminococcus, and Prevotella*) play a pivotal role in the assembly of the two core Ruminosignatures identified in our study. This suggests that their influence on community structure is both disproportionate and mechanistically grounded in their metabolic versatility. Such generalism behaviour likely contributes to RS assembly and to the metabolic support of non-core taxa, reinforcing their central role in maintaining rumen functional stability, community organization, and host–microbiome symbiosis. Together, these observations suggest that the host-associated patterns captured by RS may, at least in part, stem from the characterized metabolic roles of core taxa described previously [46], reinforcing the view that their contribution to host metabolism is an emergent property of coordinated microbial behavior rather than isolated taxon-specific effects.

To identify microbial CAGs associated with CH₄ emissions, we tested associations between individual Ruminosignatures abundances and CH₄ output, revealing multiple RS linked to methane emissions. However, the direction and strength of this link vary across cohorts (Figure 5, Supplementary Figure 11). Among them, despite substantial variation in environmental conditions, farm management practices, host genetic background, and CH₄ measurement methodologies, the abundance of the RS-UBA2 was consistently negatively associated with methane emissions across the full dataset, as well as with the acetate:propionate ratio in the Canadian cohort. Moreover, a positive link of RS-UBA2 with feed efficiency was observed in the Italian Holstein cows and the Irish crossbred populations. The genus *UBA2810* that drives RS-UBA2, was classified within the Gammaproteobacteria and exhibits phylogenetic affinity to the family *Succinivibrionaceae*. Members of this lineage are known for their ability to produce succinate, primarily through the reduction of fumarate. Succinate production, similar to propionate formation, constitutes a hydrogenotrophic process that can compete directly with methanogens for available hydrogen, thereby limiting methane formation [47]. In fact, several studies have consistently reported an association between members of the Succinivibrionaceae family and CH_4_ production in ruminants and other herbivores [18, 19, 48, 49, 50]. The association of these bacteria with lower methane emissions may be attributed to their hydrogenotrophic metabolism, which limits availability of hydrogen, an essential electron donor in methanogenesis. In agreement with this proposed mechanism, a negative correlation between the abundance of RS-UBA2 and the abundance of the methanogenic archaea *Methanobrevibacter* was observed across all populations (R_Holstein_ = -0.39; R_Canada_ = -0.33; R_Ireland_ = -0.25). Together, these patterns suggest that the host-associated effects captured by this Ruminosignature likely reflect the previously characterized metabolic role of Succinivibrionaceae in hydrogen redirection, reinforcing their contribution to methane mitigation at the level of coordinated microbial behavior, thereby strengthening the interpretation that Ruminosignatures capture metabolically grounded, functionally coordinated microbial behavior linked to host phenotypes. Additionally, in the Holstein discovery population, the abundance of RS-UBA2 showed strong negative correlation (R = -0.52, P < 0.001) with the abundance of the holotrich protozoa *Dasytricha*, but displayed a positive relationship with the abundance of *Entodinium* (R = 0.51, P < 0.001). *Methanobrevibacter* is the dominant genus of rumen methanogens playing a major role in CH_4_ production via hydrogenotrophic metabolism. *Dasytricha* produces large amounts of hydrogen, which *Methanobrevibacter* use as substrate to generate CH_4_ [51]. In fact, a positive association of *Methanobrevibacter* and *Dasytricha* with CH_4_ and the symbiotic relationship between them is well documented (18, 52, 53, 54).

In agreement with previous studies [11], the negative association between CH₄ emissions and the abundance of RS-*Prev* was not consistent across populations, which may reflect functional heterogeneity within the *Prevotella* genus. *Prevotella*, a metabolically versatile taxon capable of processing a wide range of proteins and polysaccharides, and one that has been repeatedly associated with low methane emissions in previous studies [46, 55, 56]. *Prevotella* is a well-characterized propionate producer [55], and propionate formation represents a metabolic route in which net hydrogen is consumed rather than released [57]. In addition, *Prevotella* is predicted to produce amino acids such as lysine and cysteine, which can serve as precursors for butyrate and propionate synthesis [46, 57]. Together, these metabolic traits suggest that RS_Prev has the capacity to serve as an alternative and channeling reducing equivalents into alternative fermentation end products. Taken together, these findings demonstrate a coherent association between the increased abundance of RS-UBA2 and context specify for RS-Prev, their predicted hydrogen-consuming metabolic functions, and reduced methane emissions. This convergence of taxonomic identity, functional potential, and phenotypic outcome supports a model in which specific bacterial Ruminosignatures actively redirect electrons flow away from methanogenesis, contributing to lower methane production in the rumen ecosystem.

However, there are some limitations in this study. The dynamic nature of microbial CAGs which shift with host age has been documented in human and pig gut microbial ecosystems [13, 15]. The current study focused exclusively on adult animals; it overlooked the age-related dynamics of microbial CAGs assembly and potentially missed critical insights into early-life development. This is particularly relevant for putative dietary interventions to modulate RS-abundance given that early life has been proposed as a window of opportunity for modulating the ruminal microbiota [58, 59, 60]. Moreover, these findings are based solely on metataxonomic information, which limits the ability to draw conclusions about the functional roles of RS. Therefore, to capture active microbial processes and gain a more comprehensive understanding, future studies should incorporate functional approaches from other source of information such as metatranscriptomics and metaproteomics.

Despite these limitations, the findings herein offer novel insights into the assembly of cattle ruminal microbiota. This study demonstrated that RS are heritable and associated with key traits in cattle. Notably, RS-UBA2 exhibited sufficient stability across multiple breeds and different environments to be considered a promising candidate proxy for potential integration into breeding programs aimed at mitigating CH_4_ emissions. Such proxies could reduce the need for long and costly CH_4_ measurement and potentially enhance the accuracy of selection when combined with host genetic information. Microbial traits can be integrated into genetic evaluations either as additional traits that are genetically correlated with host traits, improving prediction accuracy, or by using microbial profiles to construct relationship matrices that explicitly capture host–microbiome interactions [7]. Overall, the implementation of NMF to infer microbial CAGs represents a step forward toward ecosystem-based modeling of the rumen microbiome, supporting the idea that ecosystem-level interactions, rather than individual microorganisms, underlie key ruminal processes influencing host productivity and environmental impact. This approach facilitates the integration of microbial community structure with host and environmental variables, opening avenues for predictive modeling, causal inference, the discovery and designing synthetic consortia for targeted microbial modulation in livestock.

## Conclusions

This study provides new insights into the assembly of the cattle rumen microbiota by introducing the Ruminosignatures concept. Commons and population-specific heritable microbial CAGs associated with CH₄ emissions and feed efficiency were identified. These findings suggest that classifying rumen microbiota into RS could play a key role in microbiome-informed precision farming and breeding programs aimed at reducing the cost of feeding animals and lowering the environmental footprint of cattle production. Yet, there are particularities associated with the breed, diet and production system that emphasize the importance of tailoring modulatory strategies to specific breeding and production systems.

## ABBREVIATIONS

NMF: Non-negative Matrix Factorization
RKHS: Reproducing Kernel Hilbert Space
GRM: Genetic Relationship Matrix
MCMC: Monte Carlo Markov Chain
CH₄: Methane
CAGs: Co-abundance groups:
PCA: Principal Component Analysis
ASV: Amplicon Sequence Variant
ECM: Energy-Corrected Milk
DMI: Dry Matter Intake
RS: Ruminosignatures
UBA2: UBA2810
Prev: Prevotella
Fibr: Fibrobacter
Trep: Treponema
Soda: Sodaphilus
Buty: Butyrivibrio_A_168226
Cryp: Cryptobacteroides
Rumi: Ruminococcus
Shar: Sharpea
ADG: Average Daily Gain
FCR: Feed Conversion Ratio
RFI: Residual feed intake
Ace: Prop: Acetate-to-Propionate ratio
IT: Italy
UK: United Kingdom
FR: France
ANG: Angus
CHAR: Charolais
HYB: Kinsella Composite Hybrid

## DECLARATIONS

### Data availability

The raw sequencing data employed in this article has been submitted to the NCBI’s Sequence Read Archive (https://www.ncbi.nlm.nih.gov/sra); and BioProject under accession number: PRJNA517480; PRJNA797238; PRJNA393057; PRJEB98694.

### Authors’ contributions

YRC and ITV conceived and designed the study, performed the data analysis and drafted the manuscript. OF and IM contributed to the data analysis, experimental design and provided key microbiome expertise. IT, PBP and DPM performed data collection, contributed to the study designed and drafted the manuscript. LLG, SMW, DK, PS, SFK, DK, DK, and RE, PAA, performed data collection, sample processing, edited the manuscript. contributed to animal trials and phenotypic data acquisition. AR and RQ provided statistical guidance and critical input on the analysis. PCG and PB contributed to the manuscript editing. DPM and YRC contributed to data interpretation and supervised the study. All authors critically reviewed, edited, and approved the final manuscript.

## Acknowledgments

We are grateful to the following people for their contributions to production of the discovery dataset from UK and Italy under the RuminOmics project: N. Saunders, S. L. Potterton, J. Craigon, E. Gregson, R. Goodman, R. H. Wilcox, L. J. Tennant, E. M. Homer, D. Li, K. Lawson, L. Silvester, G. Fielding-Martin, N. F. Meades, L. Billsborrow, N. Armstrong, I. Norkiene, and S. Northover (University of Nottingham, UK); A. Minuti, E. Trevisi, M. L. Callegari, F. P. Cappelli (Università Cattolica del Sacro Cuore, Italy); K. O. Fliegerová, J. Mrázek, H. Sechovcová, J. Kopečný (CAS, Czech Republic); A. Bonin, F. Boyer, P. Taberlet (Universitaire de St Martin d’Hères, France); F. Strozzi, F. Biscarini, J. L. Williams (Parco Tecnologico Padano, Italy); R. J. Wallace (University of Aberdeen, UK); K. J. Shingfield (Luke, Finland). The authors also warmly thank all technical staff at the INRA Méjusseaume farm, and Gilles Renand (INRAe, France) for his valuable support.

